# kRISP-meR: A Reference-free Guide-RNA Design Tool for CRISPR/Cas9

**DOI:** 10.1101/869115

**Authors:** Mahmudur Rahman Hera, Wahid Al Azad Navid, Fuad Ahmed Udoy, Amatur Rahman, Atif Rahman

## Abstract

Designing specific guide RNAs for CRISPR/Cas9 typically relies on reference genomes, which limits the applicability in non-model organisms and fails to account for individual-specific genomic variations. To address this, we present a reference-free method for single-guide RNA (sgRNA) design that operates directly on raw sequencing reads, implemented in a tool called kRISP-meR. Instead of relying on whole genome assembly, kRISP-meR models the ***`k`***-mer frequency spectrum of the sequencing data using an Expectation-Maximization (EM) algorithm and uses a probabilistic model to calculate the expected number of cuts for candidate gRNAs by estimating genomic copy numbers. Results on datasets from three different organisms show that the expected cuts estimated by kRISP-meR from reads agree closely with those computed from reference assemblies. Furthermore, analysis of human and rice sequencing data reveals that kRISP-meR successfully detects and filters candidate gRNAs within repetitive genomic regions with high off-target risk, highlighting its importance for gRNA design when references are unavailable or incomplete.

## 1 Introduction

The CRISPR/Cas9 system is a groundbreaking technique for genome-editing that promises precise modification of target DNA sequences in diverse organisms [1–3]. In this system, a guide RNA (gRNA) directs the Cas9 nuclease to specific genomic loci, where it induces double-strand breaks that are subsequently repaired, enabling gene disruption or targeted sequence modification. These capabilities have led to its broad adoption in functional genomics, vector control, and therapeutic genome editing, including recent clinical demonstrations of in vivo and ex vivo gene editing [4, 5].

A critical determinant of the CRISPR/Cas9 performance is the design of sgRNAs that achieve high on-target activity while minimizing off-target effects. Off-target editing, resulting from partial sequence homology at unintended genomic loci, is a major concern due to its potential to introduce harmful mutations and compromise genomic integrity [6, 7]. To address this challenge, numerous computational tools have been developed to predict sgRNA efficiency and specificity, including widely used frameworks such as GuideScan [8–10] and CRISPOR [11].

On-target activity scores estimate how effectively a guide produces a cut at its intended target, whereas off-target scoring methods estimate the likelihood of cleavage at unintended genomic sites. Doench et al. introduced Rule Set 2 (RS2), trained on pooled screen data to predict sgRNA activity, and the cutting frequency determination (CFD) score to estimate the effects of mismatches on off-target cleavage [12]. In the same study, both scores were used together to design the optimized human and mouse sgRNA libraries Brunello and Brie. Separately, Haeussler et al. evaluated multiple on-target and off-target scoring methods and integrated them into CRISPOR to facilitate guide selection [13].

Despite substantial progress, most existing approaches design sgRNAs using a reference genome. This process is increasingly inconsistent with current understanding of genomic diversity. High-quality genome assemblies, although improving, remain unavailable for many non-model organisms, limiting the applicability of reference-based design tools [14]. Even in well-characterized species such as humans, recent advances in long-read sequencing and telomere-to-telomere assembly have revealed substantial previously unresolved regions of the genome [15]. Furthermore, pangenome analyses have demonstrated that a single reference genome fails to capture the full spectrum of structural variation across individuals, including insertions, deletions, and segmental duplications that may introduce additional off-target sites [16, 17].

These limitations have direct implications for the design of sgRNA. When off-target prediction is performed against an incomplete or non-representative reference, homologous genomic regions may be missed, leading to overestimation of sgRNA specificity. In practice, this can result in the selection of guides that appear unique in the reference genome but cause unintended binding in the target individual, leading to off-target double-stranded breaks and edits.

Several recent methods have sought to incorporate individual-specific variation into sgRNA design by integrating sequencing data with reference genomes. Tools such as AlleleAnalyzer, CrisFlash, and CRISPR-HAWK account for single-nucleotide variants and short insertions or deletions through variant-aware alignment [18–20]. However, these approaches remain fundamentally reference-dependent and are therefore limited in their ability to capture larger structural variations and duplicated regions that are not represented in the reference assembly.

Here, we present kRISP-meR, a reference-free computational framework for sgRNA design that operates directly on raw sequencing reads from the target individual. Given a user-defined target region, kRISP-meR decomposes sequencing reads into *k*-mers and employs an Expectation-Maximization (EM) [21] algorithm-based probabilistic framework to estimate genomic copy number and evaluate off-target potential without requiring genome assembly or alignment. We show that kRISP-meR recovers most high-quality guides identified by established reference-based tools such as GuideScan2 [10] and CRISPOR [11]. Crucially, we demonstrate that kRISP-meR outperforms these methods by successfully identifying and eliminating high-risk guides that target loci appearing as single-copy in the reference assembly, but which are actually duplicated in the target individual’s genome.

## 2 Results

### 2.1 Overview of kRISP-meR

An overview of the methodology of kRISP-meR is illustrated in Figure 1 with each step described in detail in the Methods section. Given a user-defined target region as a sequence and sequencing reads from the individual, kRISP-meR first aligns the reads to the target sequence using Bowtie 2 [22] and refines the target sequence with Pilon [23] to capture sample-specific variation. This step can be skipped if the target sequence is known precisely. Candidate gRNAs are then identified from the resulting personalized target sequence at each compatible protospacer-adjacent motif (PAM), NGG by default.

**Fig. 1:**
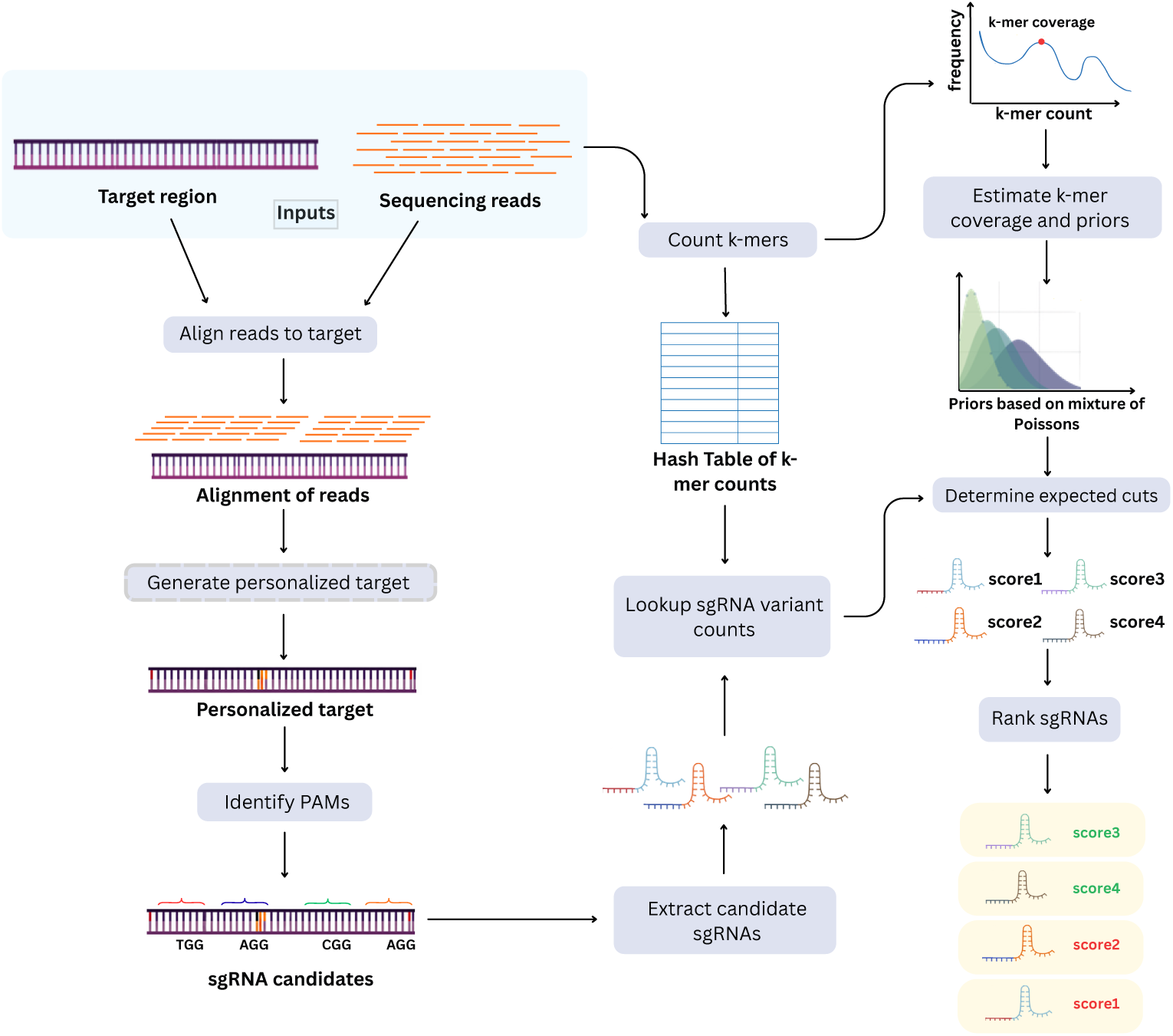
Overview of kRISP-meR. kRISP-meR takes as input a target sequence and sequenced reads from an individual. It starts by aligning the reads to the target sequence to detect sample specific variations and to generate a personalized target. Then the candidate guide RNAs are identified. Next, the number of times *k*-mers appear in the sequencing reads are determined, and *k*-mer coverage is estimated using the EM algorithm. Finally, scores to indicate efficiency and specificity of each candidate gRNA are determined using the *k*-mer counts with the use of probabilistic models.

Next, kRISP-meR counts canonical *k*-mers in the reads using Jellyfish 2 [24]. We set *k* = 23, so that one *k*-mer spans the 20 nt protospacer and its PAM. The resulting *k*-mer spectrum is modeled as a mixture of Poisson distributions. From this model, kRISP-meR infers the *k*-mer coverage and a prior over genomic copy number. The parameters are learned using the expectation–maximization (EM) algorithm [21]. For any *k*-mer, this gives a posterior distribution over its genomic copy number, given how often it was seen in the reads.

For each candidate guide, kRISP-meR then calculates a score representing the expected number of cuts by that guide across the entire genome (Scoring Candidate gRNAs in Methods). It first enumerates every *k*-mer within Hamming distance *d* of the guide’s *k*-mer (*d* = 2 by default) and stores this set in a trie to remove duplicates. The read count of each of these *k*-mers is then retrieved by querying the Jellyfish hash table directly. For each *k*-mer found in the reads, we multiply its expected genomic copy number by its CFD score based on the cutting frequency determination model [12]. The expected number of cuts is the sum of these products.

In addition to the expected number of cuts, kRISP-meR reports the following quantities for each guide. The first is an estimate of how many times the target region itself occurs in the genome, obtained by selecting the copy number that best explains the *k*-mer spectrum of the target alone. The second is the expected number of cuts within the target region. An inverse-specificity score combining the three is also reported, defined in Additional Quantities for gRNAs in Methods. Finally, an on-target activity score based on the Rule Set 2 [12], computed from the 30 bp genomic context of the guide, is also provided. The output also records the GC content and the strand on which each guide lies.

kRISP-meR ranks the guides based on the expected number of cuts from the lowest to the highest and reports the above mentioned quantities for all the guides. However, we recommend using guides with an expected number of cuts below 1.5 since it indicates the guide is likely to cut the genome once and all the analysis reported in this paper is based on this threshold.

### 2.2 Benchmarking Against Reference-Based Tools

To evaluate the performance and reliability of kRISP-meR, we benchmarked it against two established reference-based tools: CRISPOR [11] and GuideScan2 [10]. For this comparative analysis, we used datasets from three organisms of varying genomic complexity: *Saccharomyces cerevisiae* (S288C [25]), *Mus musculus* (GRCm39 [26]), and *Homo sapiens* (GRCh38.p13 [27]). The three species were chosen to ensure the availability of reliable reference genomes for accurate comparison with reference-based tools as well as real whole-genome sequencing data (Supplementary Table 1), so that the *k*-mer spectra reflect genuine error profiles, coverage variation, and library biases. For each organism, random target sequences were systematically generated across three distinct genomic contexts: protein-coding regions, non-coding regions, and duplicated regions following the approach in [8], with 50 targets drawn from each context, giving 150 target regions per organism.

First, we assess the probabilistic model of kRISP-meR by comparing the expected number of cuts predicted by kRISP-meR from raw sequencing reads with the baseline scores computed using the reference genomes. The expected number of cuts calculated based on reads versus the scores based on references, i.e., assemblies for the three species are shown in Figures 2(a-c). We find that read-derived and reference-derived estimates largely agree, with points scattered mostly along the diagonal (Pearson *r* = 0.97, 0.99 and 0.99 for *S. cerevisiae*, *M. musculus* and *H. sapiens* respectively; Spearman *ρ* = 0.94, 0.99 and 0.78; all *p <* 0.001; *n* = 1585, 1779 and 2069 guides). The comparatively low Spearman correlation of 0.78 in *H. sapiens* is driven by many guides sharing the same expected number of cuts of 1 with slightly different read-based scores, which reduces rank-based correlation without suggesting a loss of linear agreement. We observe that the read-based scores calculated by kRISP-meR are sometimes slightly higher than the reference-based scores, indicating that kRISP-meR is conservative in its selection of gRNA while the occasional substantial differences between the two scores are possibly due to repeats collapsed in the assembly that are captured in the reads.

**Fig. 2:**
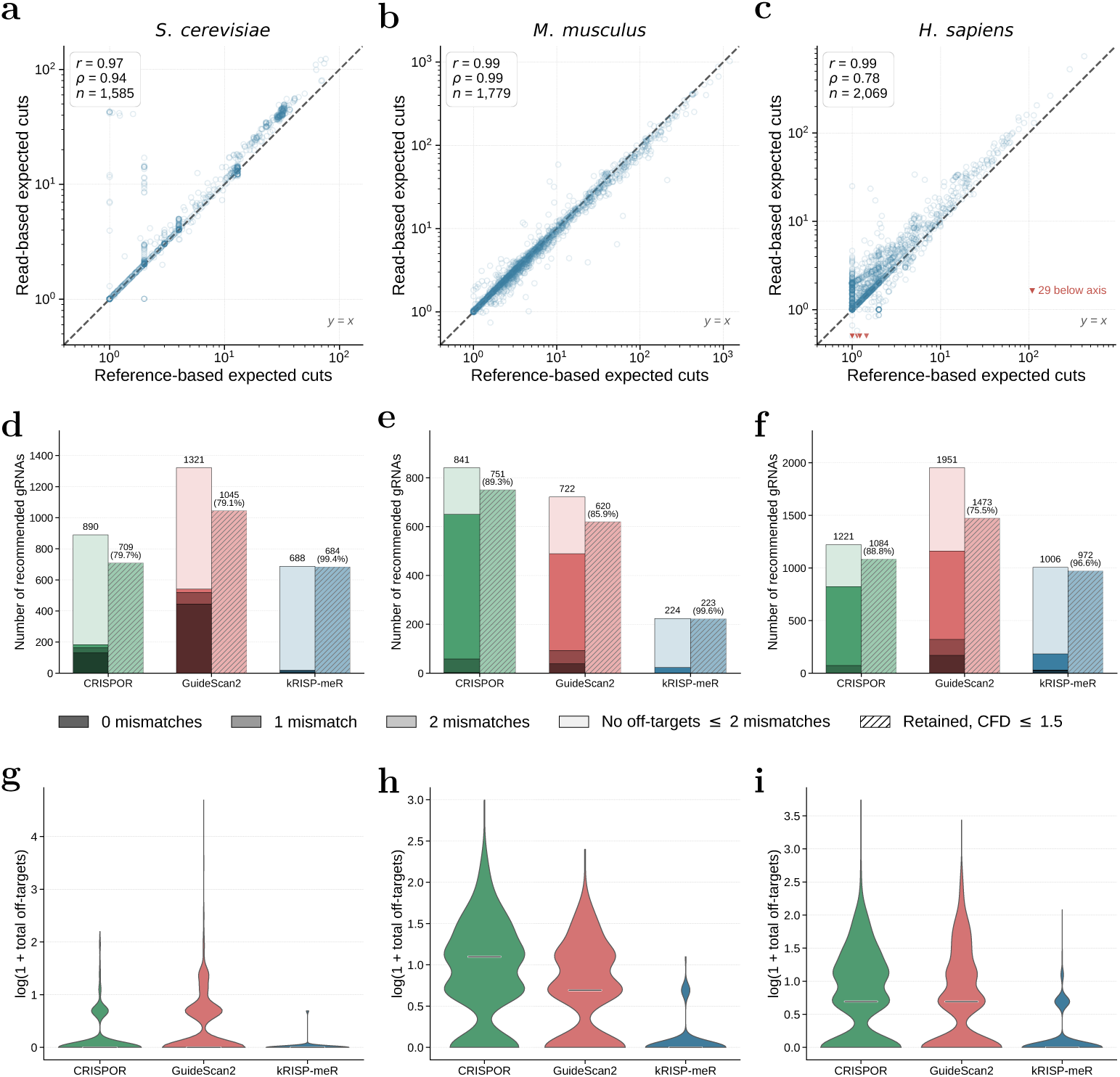
Comparison of kRISP-meR with GuideScan2 and CRISPOR across three genomes (*S. cerevisiae*, *M. musculus*, and *H. sapiens*). **(a–c)** Scatter plots showing the relationship between the expected cuts calculated via the complete reference assembly (x-axis) and the read-based estimations produced by kRISP-meR (y-axis). **(d–f)** Comparison of off-target activity profiles for the candidate sgRNAs recommended by each tool. The solid bars categorize guides by their most severe off-target occurrence (0, 1, or 2 mismatches, or no off-target site at any distance considered); the bar total is the size of each tool’s recommended set. The adjacent hatched bars give the number of those same guides retained after a strict safety filter using the cutting frequency determination (CFD) pipeline (expected cuts *≤* 1.5), with the retention percentage computed over the same guide set. **(g–i)** Distribution of the total number of off-target sites per recommended guide, summed over 0–2 mismatches without weighting and plotted as ln(1 + *n*). kRISP-meR’s distribution is concentrated near zero in all three species.

Next, we evaluate the off-target effects of the guides suggested by each tool across all target regions. We focused exclusively on guides with acceptable predicted specificity and applied tool-specific filtering criteria. Candidate sgRNAs were retained only if they met the following thresholds: an expected number of cuts below 1.5 for kRISPmeR (retention at other thresholds is shown in Supplementary Figure 2 and discussed in Supplementary Note 1), a specificity score exceeding 50 for CRISPOR, and a specificity score above 0.2 for GuideScan2. These thresholds were selected to represent the standard operational guidelines recommended by the respective tools for precise editing.

Figures 2(d-f) present breakdowns of the gRNAs suggested by each tool based on their off-target effects for the three species, while the corresponding region-specific results for protein-coding, duplicated and non-coding targets are provided in Supplementary Figure 3. For each tool, the first bar categorizes the recommended gRNAs by their most severe off-target occurrence (0, 1 and 2 mismatches). The reference-based methods, particularly GuideScan2, proposed a substantial number of guides with off-target sites with 0-mismatch (perfect off-target) and 1-mismatch in the actual genome. In contrast, kRISP-meR’s utilization of raw read *k*-mer spectra inherently flags and removes these high-risk candidates. As a result, its gRNA pool is composed almost entirely of guides with off-target sites with 2 mismatches. The underlying counts are given in Supplementary Table 2.

We also passed every generated gRNA from all three tools through the CFD scoring pipeline [12] on the reference genomes, applying a filter to retain only guides with expected cuts of 1.5 or less. The second bar in Figures 2(d-f) for each tool illustrates the retention rate after filtering, computed over the same recommended guide sets. kRISP-meR retained 99.4% of its guides in the yeast dataset, 99.6% in the mouse dataset, and 96.6% in the human dataset, which is higher than the reference-based tools. CRISPOR retained between 79.7% and 89.3% of its candidates, and GuideScan2 between 75.5% and 85.9%, across the three species. It is worth noting that the selection rule used by kRISP-meR and the downstream CFD-based validation filter are based on the same quantity: a CFD-weighted sum of expected cuts. Therefore, higher retention rates of kRISP-meR are expected. However, there are fundamental differences in the two calculations. The validation filter uses copy-number information taken directly from the reference assembly which is available to the reference-based tools, whereas kRISP-meR estimates copy numbers from raw-read *k*-mer abundance without using the reference genomes.

To further quantify the global specificity of the candidate sgRNAs and assess the cumulative risk of each guide in the generated set, we examined the overall distribution of the off-target scores for every recommended guide. Figures 2(g-i) show, for each tool, the distribution of the total number of off-target sites per recommended guide. For each guide, the sites at 0, 1 and 2 mismatches are added into a single count *n*, without weighting, and plotted as ln(1 + *n*). A logarithmic scale was utilized to effectively compress highly skewed distributions, so that guides with almost no off-target sites remain distinguishable from those with hundreds.

The distributions for both reference-based tools, CRISPOR and GuideScan2, exhibit wide variances, and prominent, elongated upper tails. These extended tails indicate that a substantial proportion of the guides recommended by these tools still carry a high cumulative number of off-target matches scattered across the genome. Even where such guides pass standard filtering criteria by avoiding 0-mismatch or 1-mismatch sites, a large number of partial matches may sequester the Cas9–RNA complex, which has been proposed to increase the amount of Cas9 required for efficient editing [28, 29]. In contrast, the off-target distribution for kRISP-meR is heavily anchored near zero, producing a highly constrained profile. This pattern is consistent across the varying genomic complexities of the human, mouse, and yeast datasets. Because kRISP-meR evaluates target abundance directly from raw sequencing reads via *k*-mer frequency spectra, it inherently penalizes guides associated with repetitive or duplicated elements that are often collapsed or misrepresented in standard reference assemblies. This indicates that kRISP-meR reduces not only the most severe off-target activity but also the global off-target footprint per guide.

### 2.3 Guide Recommendations Under Individual–Reference Genomic Discordance

We now turn to comparing the performance of kRISP-meR with that of reference-based tools in situations where the genomes of individuals differ from the reference genomes. To evaluate whether guide recommendations reflect the genomic context of the target individual rather than that of a standard reference assembly, we used the CHM13 human assembly (*Homo sapiens*) [30] and the N22 rice assembly (*Oryza sativa*)[31] as operational ground-truth representations of the individual specific genomes. The corresponding standard reference assemblies used were GRCh38.p13 for human (*H. sapiens*) and the Nipponbare japonica assembly for rice (*O. sativa*) [32]. The sequencing datasets from CHM13 and N22 used in this analysis as well as the assembly accessions are listed in Supplementary Table 3.

We constructed four categories of sequences where the individual genomes and the corresponding reference assemblies disagree: (i) sequences with multiple-copies in the individual but single-copies in the reference, (ii) sequences with single-copies in the individual but multiple-copies in the reference, (iii) sequences present in the individual but absent from the reference, and (iv) sequences absent from the individual but present in the reference. These categories were defined at the *k*-mer level by comparing the copy number of each guide *k*-mer between the individual and reference assemblies. The genome-wide distributions of candidate guide *k*-mers across these categories are shown in Figures 3(a–d). We find that as much as 24.4% of the *k*-mers corresponding to candidate gRNAs in the *Oryza sativa* reference are absent from the N22 assembly while 1.6% are present more than once. Similarly, 2.3% and 1.0% of the candidate guides in the *Homo sapiens* reference do not appear and appear more than once in the CHM13 assembly, respectively.

**Fig. 3:**
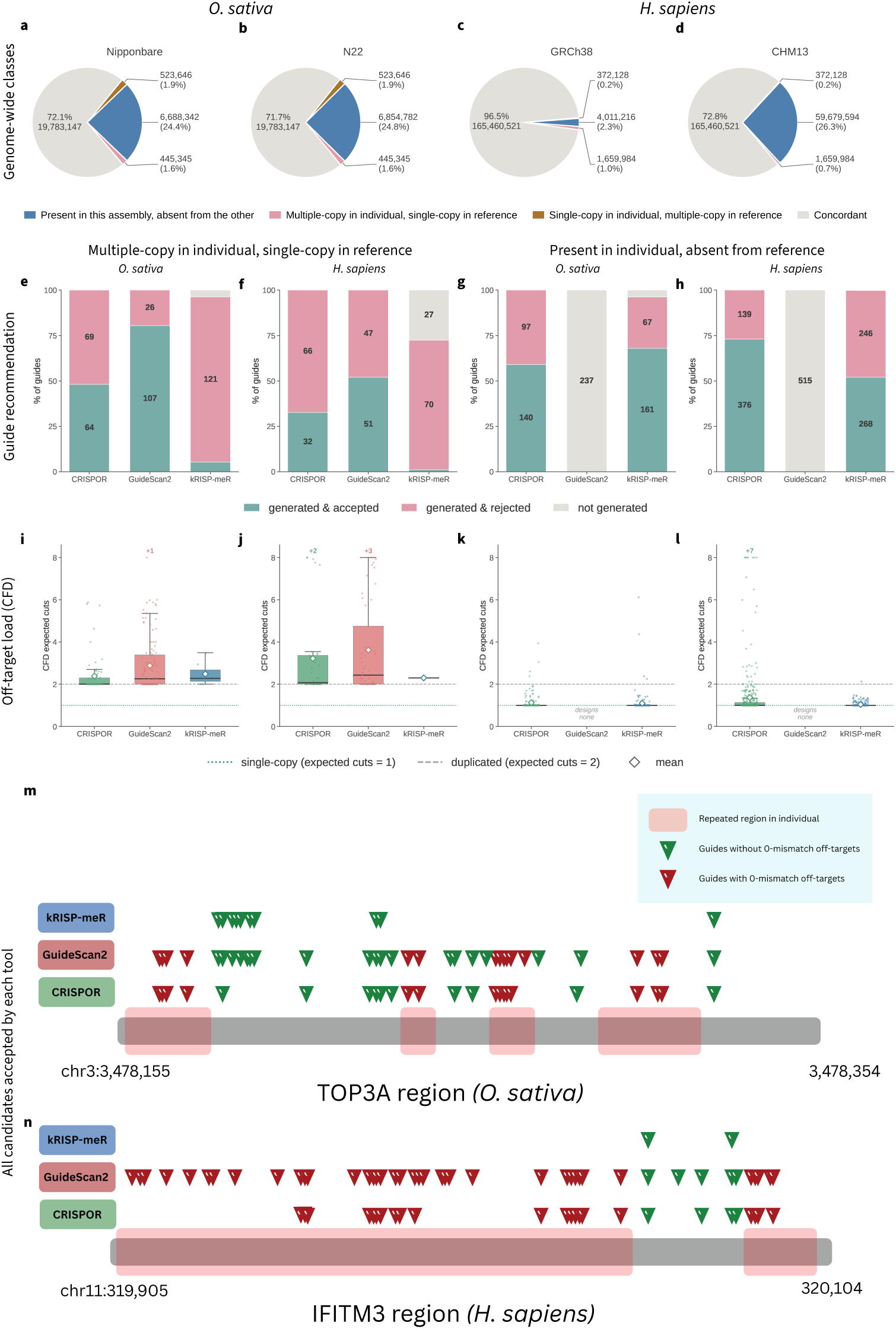
Genome-wide class composition, guide recommendations, and off-target load under individual–reference genomic discordance in rice (*O. sativa*; N22 versus japonica) and human (*H. sapiens*; CHM^9^13 versus GRCh38). **(a–d)** Genome-wide distribution of candidate guide *k*-mers across four discordance categories and the concordant remainder, shown separately for each assembly. **(e–h)** Class-defining guides that each tool accepted, rejected, or did not generate, shown as a percentage of the class with counts labelled, at loci multiple-copy in the individual but single-copy in the reference (e, f) and loci present in the individual but absent from the reference (g, h); guides should be rejected in the former and recovered in the latter. **(i–l)** Assembly-CFD expected genomic cuts for the guides each tool accepted, in the corresponding panels; dotted and dashed lines denote one and two expected cuts, and the diamond marks the mean. **(m, n)** Guides at *TOP3A* (chr3:3,478,155–3,478,354, OsN22_03G004820) and *IFITM3* (chr11:319,905–320,104, CHM13), 200 bp each. Triangles mark the positions of guides each tool accepted. Pink denotes a region repeated in the individual; green triangles have no 0-mismatch off-target in the individual’s assembly, red at least one.

We then extracted a 150 bp target window centered on 50 randomly selected guides from each class, and the guides within these windows constituted the candidate pool against which the output of each tool was evaluated. It is worth stressing that the individual assemblies were used only to define genomic discordance categories and the ground-truth candidate pool, and subsequently for estimation of off-target effects. They were not provided as input to any of the tools. Instead, kRISP-meR was run on raw sequencing reads from the individuals. In contrast, CRISPOR and GuideScan2 were run using the corresponding standard reference assemblies, as required by their reference-dependent workflows. In the main text, we focus on two discordance classes with direct implications for gRNA safety and practical experimental design: sequences that are single-copy in the reference but multiple-copy in the individual, and sequences that are present in the individual but absent from the reference. Results for the two complementary categories are provided in Supplementary Figure 4.

We first considered targets that are single-copy in the reference but multiple-copy in the individual. Guides targeting such loci should ideally be rejected or strongly down-weighted because the repeats in the individual increases the potential for unintended edits. As shown in Figures 3(e, f), CRISPOR and GuideScan2 generated and accepted a large fraction of guides in this category because both tools score specificity using the reference, where the locus appears single-copy. In contrast, kRISP-meR rejected or did not generate majority of these guides because its copy number estimate is derived from the individual’s sequencing reads and therefore reflects the duplications. It accepted 7 of 133 guides from this class in rice and 1 of 98 in human, against 64 and 32 for CRISPOR and 107 and 51 for GuideScan2. The expected-cut distributions of the accepted guides are shown in Figures 3(i, j). We observe that although the medians are comparable, the reference-based tools accept guides reaching 5.88 and 8.86 expected cuts, and up to 34.8 in human, whereas no guide accepted by kRISP-meR exceeds 3.49.

We next evaluated targets that are present in the individual but absent from the reference. Such loci represent valid editing targets in the individual, and an effective guide-design tool should be able to identify and evaluate guides for them. However, we find that reference-dependent tools are structurally limited in their ability to assess these targets as the underlying sequence is absent from the reference assembly. As shown in Figures 3(g, h), GuideScan2 and CRISPOR exhibited this limitation in different ways. GuideScan2 did not generate any guide from targets in this category, since it enumerates candidates from a precomputed reference-genome index that does not contain the target sequence. CRISPOR accepts a user-provided target sequence, so it generated candidates and accepted a large fraction of them: 140 of 237 in rice and 376 of 515 in human. However, the specificity scores it reports are inaccurate since they are based on the references. In contrast, kRISP-meR generated and evaluated guides using sequencing reads from the individual, in which the target sequence is present. It accepted 161 of 237 guides in rice and 268 of 515 in human, rejecting those with read-derived evidence of additional cutting sites. The expected-cut distributions of the accepted guides in Figures 3(k, l) show the consequence of scoring using the wrong genome. Both tools center on a median of 1.00, as they should at a single-copy locus, but 7.7% of the guides CRISPOR accepts in human exceed two expected cuts, reaching 11.9, against 0.4% and a maximum of 2.13 for kRISP-meR.

Our analysis also revealed that many of the candidate guides from various discordant classes are located within protein-coding regions. Intersecting protein-coding regions with the multiple-copy in individual and single-copy in the reference and the present in individual and absent from the reference classes yielded 10,713 target regions in *O. sativa* and 583 in *H. sapiens*, with one target window per gene. From these, we examined two loci belonging to closely related paralogous gene families: *IFITM3* in *H. sapiens*, an interferon induced restriction factor that lies within a tandem cluster of *IFITM* paralogues, [33] and *TOP3A* in *O. sativa*, a type IA DNA topoisomerase whose paralogue *TOP3B* arose by an antient gene duplication [34]. Paralogous sequences present a particular challenge for reference-based off-target assessment because the duplicated regions represented in a reference assembly may not fully reflect the copy number and sequence composition of an individual.

The guides generated by kRISP-meR, GuideScan2 and CRISPOR at these loci are summarized in Figures 3(m, n) (23-mer profiles of the target regions are shown in Supplementary Figures 5 and 6). For the target region in *TOP3A*, GuideScan2 accepted 35 of 36 candidates, 15 with 0-mismatch off-target sequences, while CRISPOR accepted 24 of 36 candidates, 14 of which had exact off-targets. kRISP-meR retained 12 of its 31 candidates, none with exact off-target sequence matches. In particular, 11 of the 12 guides retained by kRISP-meR were also identified by other tools, but kRISPmeR excluded candidates with predicted off-targets under its selection criterion. For *IFITM3*, GuideScan2 generated 46 candidates and accepted all of them, 41 with at least one exact off-target site, while CRISPOR accepted 25 of 46 candidates, 21 of which had 0-mismatch off-targets. kRISP-meR retained 2 of its 22 candidates, none with exact off-target. This illustrates how kRISP-meR’s read-based estimation can exclude potentially unsafe guides with predicted off-targets that are accepted by reference-based tools. We also ran all three tools on the full *TOP3A* gene (10,145 bp) and the *IFITM3* gene (7,709 bp) as target regions. kRISP-meR accepted a smaller proportion of guides carrying a perfect off-target site than CRISPOR or GuideScan2 at both loci (Supplementary Table 4).

## 3 Discussion

Precise genome editing using CRISPR/Cas9 technology requires the design of guide RNAs that minimize off-target activity. Existing tools for designing gRNAs rely primarily on reference genomes of organisms. This makes them difficult to apply to non-model organisms with no or incomplete reference genomes. Furthermore, they are not well-suited for personalized genome editing as the genome of the individual may contain regions not reflected in the reference.

To overcome these challenges, we present a reference-free method for sgRNA design implemented in a tool called kRISP-meR. It takes as input a target region and sequencing reads from the target individual, and estimates the off-target effects of the candidate guides in the target region based on *k*-mer counts in the reads. Our results show that genomic copy numbers can be inferred directly from sequencing reads and used to assess sgRNA specificity without a reference genome. Across human, mouse and yeast, the expected number of cuts estimated from reads agreed closely with estimates computed from the references.

We also compare the performance of kRISP-meR and reference-based tools using datasets from a human individual (CHM13) and a rice cultivar (N22) whose genomes deviate from references. We focus on guides in regions with differing numbers of copies in the references and in the individuals. We find that reference-based tools largely accepted guides that appear single-copy in the references GRCh38 or Nipponbare but occur at several loci in CHM13 or N22, while kRISP-meR detected the extra copies using the reads and rejected the guides. At the paralogous loci *IFITM3* in human and *TOP3A* in rice genomes, kRISP-meR only retained guides without exact off-target sites, while reference-based tools accepted guides with identical off-target sequences. Sequences present in the individual but absent from the reference expose a second weakness of reference-based design. They either cannot generate candidates for them because of their reliance on the reference, or score the specificity of the guides incorrectly when the sequence is missing in the reference. kRISP-meR uses reads from the individuals, so it evaluates such targets in their actual genomic context. Thus, it can recover individual-specific editing targets that are unavailable to reference-based tools.

kRISP-meR thus can design guides without a reference and identify a class of off-target risks that reference-based tools miss. However, several future directions remain. First, *k*-mer counts show that a sequence occurs more than once, but not where. kRISP-meR therefore cannot tell whether an off-target site falls in a coding region. It is also currently unable to incorporate a broader sequence context in its off-target effects model. Second, off-target sites are searched up to Hamming distance 2. In the CFD model, cutting probabilities decrease steeply with mismatches, but we have not measured the contribution of more distant sites. Finally, the discordance analysis relied on assemblies as ground truths that may themselves contain copy number errors.

In summary, kRISP-meR designs guides from the reads of the individual being edited, without reference genomes or assemblies. Although it is not a replacement for reference-based tools, it adds an individual-specific check to the conventional guide design approach. It is most useful when the reference is an inadequate representation of the genome to be edited, which is often the case with non-model organisms, populations divergent from the reference, and individuals or cultivars with substantial structural variation. With accurate whole genome assembly remaining a challenge, a reference-free tool such as kRISP-meR enables gRNA design for accurate and safe personalized genome editing and editing in non-model organisms that reference-based tools cannot address.

## 4 Methods

### 4.1 Identification of Candidate gRNAs

kRISP-meR takes as input sequencing reads from the individual whose genome is to be edited and a target region representing where the editing is intended. Sequencing reads are aligned to the target sequence using Bowtie 2 [22], and variants are identified using Pilon [23] to construct a personalized target sequence for the individual. If the target sequence is known exactly, this step can be skipped. The personalized sequence is then scanned for protospacer adjacent motif (PAM) sites, which are NGG by default. For each PAM occurrence, the 20 bp upstream sequence is extracted as the protospacer. Each protospacer, together with its associated PAM, forms a 23 bp candidate gRNA.

### 4.2 *k*-mer Counting from Sequencing Reads

As kRISP-meR operates without a reference genome, it relies on *k*-mer counts computed directly from sequencing reads. All *k*-mers of length *k*, corresponding to the combined length of the protospacer and PAM (*k* = 23 by default), are counted using Jellyfish 2 [24]. Counts are computed on canonical *k*-mers, so that a sequence and its reverse complement are treated as the same *k*-mer; this reflects the fact that Cas9 can cleave a site on either strand.

From these counts we construct the *k*-mer frequency histogram *H*, where *H*(*c*) is the number of distinct *k*-mers observed exactly *c* times in the read set. Throughout, *c* denotes an observed count and *i* denotes a genomic copy number. The histogram characteristically exhibits a peak at low counts arising from sequencing errors, followed by a peak at the *k*-mer coverage *λ* corresponding to single-copy genomic *k*-mers, and further peaks at approximately 2*λ,* 3*λ, …* corresponding to sequences present at higher copy numbers. Supplementary Figure 1 shows this histogram for the *H. sapiens* read set. The initial estimate of *λ* is obtained by smoothing *H* with a Savitzky–Golay filter [35] (window 9 by default) and taking the first local maximum of the smoothed curve; the preceding local minimum is recorded as the inversion point separating the error component from the genomic signal. This estimate is subsequently refined by the procedure described in Section 4.5.

The choice of *k* is determined by the biology. It must equal the protospacer plus PAM length so that a single *k*-mer count corresponds to the abundance of a complete candidate cleavage site. For PAM sequences of different length, *k* is adjusted accordingly.

### 4.3 Scoring Candidate gRNAs

kRISP-meR scores each gRNA by estimating its expected number of cuts in the genome. In CRISPR/Cas9 editing, a guide can cut not only its exact target sequence, but also similar genomic sequences containing a small number of mismatches, although such mismatched sites generally have lower cutting probability. Let K be the set of all *k*-mers within Hamming distance *d* of the gRNA target sequence, where the default is *d* = 2. The expected total number of genomic cuts is then defined as follows:

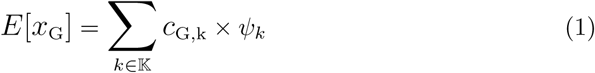

where *c*_G,k_ is the number of occurrences of *k*-mer *k* in the genome and *ψ_k_*is the probability of cleavage at that site, determined by the experimentally derived CFD mismatch matrix [12].

Since genomic copy numbers are not directly known in the absence of a reference genome, we estimate them from the *k*-mer counts observed in the sequencing reads. Specifically, we compute the expected number of cuts by taking the expectation over possible genomic copy numbers conditioned on the observed read count *c*_R,k_, using Bayes’ rule:

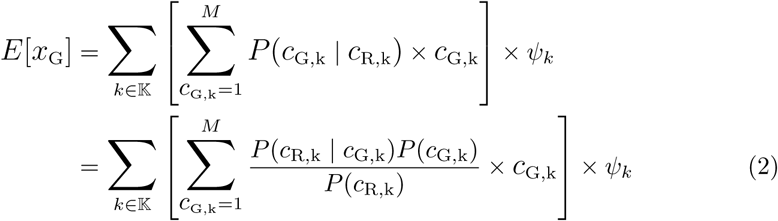

The probabilities appearing in Equation 2 are calculated as follows:

- Likelihood: Given that a *k*-mer is present *c*_G,k_ times in the genome, its count in the reads is modeled as Poisson with mean proportional to the copy number:

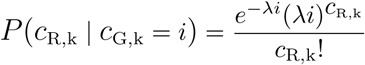

where *λ* is the *k*-mer coverage of the dataset. Poisson distributions are widely used to model *k*-mer abundance in sequencing data [36, 37].
- Prior: *P* (*c*_G,k_ = *i*) = *π_i_* is the prior probability that an arbitrary *k*-mer is present in *i* copies in the genome. The mixture weights *π_i_* are not assumed a priori but are estimated from the data as described in Section 4.5, since organisms differ substantially in repeat content.
- Marginal: The denominator is the marginal likelihood, obtained by summing the numerator over all copy numbers considered:

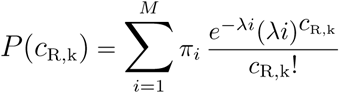

The sums in Equation 2 are truncated at a maximum copy number *M*, left at its default of 50 in all experiments. The Poisson likelihood decays rapidly once *λi* exceeds the observed count, so higher components carry negligible posterior mass, and *k*-mers genuinely present at very high copy number receive expected number of cuts far above any usable threshold regardless of *M*.

Candidate gRNAs are ranked according to their expected number of cuts in the genome, *E*[*x*_G_], with lower values indicating fewer potential genomic cuts. The gRNAs with *x*_G_ less than 1.5 are recommended since they are unlikely to cut the genome more than once (Supplementary Note 1).

### 4.4 Additional Quantities for gRNAs

In addition to the expected number of cuts in the genome *E*[*x_G_*] for every candidate gRNA, kRISP-meR reports the expected number of cuts within the target region *E*[*x_t_*], the estimated number of copies of the target region in the genome *c_G,t_*, and an inverse-specificity score that combines them. In cases where the *x*_G_ exceeds the recommended threshold, these quantities can be used to assess whether the guide is expected to cut more than once within the target region.

The value *E*[*x*_t_] is computed directly from the supplied target sequence: for each *k*-mer having PAM sequence within Hamming distance *d* of the gRNA, its occurrences in the target are counted exactly and weighted by the same CFD probability used in Equation 1. No probabilistic estimation is required, because the target sequence is given.

The target copy number *c*_G,t_ is not known, however. Although the target sequence is provided, the number of times it occurs in the genome is unknown. So it is estimated from the reads. We build the *k*-mer count histogram restricted to *k*-mers occurring in the target region, *H_t_*, and select the copy number that best explains it. A target present in *i* copies produces *k*-mer counts drawn from a Poisson with mean *iλ* (Equation 3), so we take

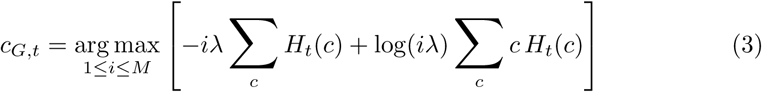

which is the maximum-likelihood copy number over the integers 1 to *M*. kRISP-meR also reports an inverse-specificity score,

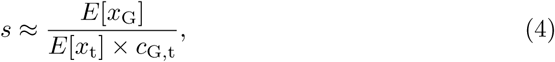

which measures how much the guide is expected to cut in the whole genome relative to the intended target site. If the target region is single-copy and the guide is expected to cut once within the target regions, the denominator is close to one and this score is nearly identical to the expected number of cuts.

Finally, kRISP-meR computes an *on-target activity* for each candidate gRNA that estimates how efficiently a guide cleaves its intended target. kRISP-meR predicts it with Rule Set 2 (RS2) [12]. The prediction uses the 30 bp sequence context of the guide, extracted from the personalized target sequence: 4 bp upstream of the protospacer, the 20 bp protospacer, the 3 bp PAM and 3 bp downstream. The resulting RS2 score is reported alongside the other guide evaluation metrics but is not used in guide ranking. Scores are computed with the Azimuth implementation of this model [12], which reproduces the published Rule Set 2 values. Guides lying within 4 bp of the start or 3 bp of the end of the supplied target do not have a complete 30 bp context and are reported without an on-target score.

### 4.5 Learning *k*-mer Coverage and Estimating Priors

To evaluate Equation 2 we require both the *k*-mer coverage *λ* and the prior distribution *{π_i_}* over genomic copy numbers. Both are estimated from the *k*-mer count histogram by fitting a finite mixture of Poisson distributions using the expectation–maximization (EM) algorithm [21].

We model the count of a genomic *k*-mer as a mixture

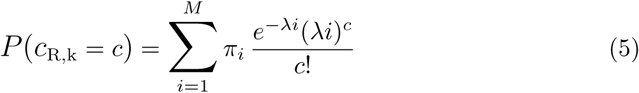

where component *i* stands for *k*-mers that occur *i* times in the genome, and 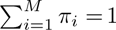. If a sequence occurs twice, reads cover it twice as often, so its counts centre on 2*λ* instead of *λ*. Every component therefore shares the same *λ*, so the means *λ,* 2*λ, …, Mλ* are fixed by one number instead of *M* separate ones. GenomeScope [37, 38] fits a similar mixture to estimate genome size, heterozygosity and repeat content. We want something narrower: the posterior over copy number for a single *k*-mer, which is what Equation 2 needs.

Sequencing errors may create *k*-mers that are not in the genome. Nearly all are seen once or twice, so the first component in the mixture is reserved for them. We found empirically that this is best modeled with a Poisson distribution initialized at a small mean of 1.01 and updated during fitting like other components. Unlike the genomic components, whose means are constrained to multiples of *λ*, the error component’s mean is free.

We use the expectation–maximization (EM) algorithm [21] to estimate the parameters. *λ* is initialized from the histogram as described in Section 4.2, and the mixture weights are initialized uniformly, *π_i_* = 1*/M* for all *i*.

**E-step:** Given the current parameters, the posterior responsibility of component *i* for a *k*-mer observed *c* times is

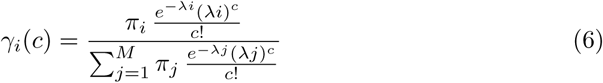

**M-step:** The mixing weights are updated as the responsibility-weighted proportion of *k*-mers assigned to each component,

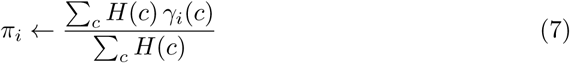

and each component mean is updated independently as the responsibility-weighted mean count,

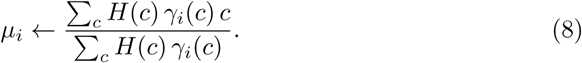

A single coverage estimate is then recovered by averaging the per-copy implied coverages *µ_i_/i* over the genomic components, weighted by the mass of each component:

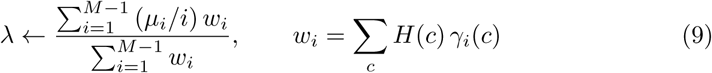

and the genomic component means are reset to *µ_i_* = *iλ* before the next iteration.

The E- and M-steps are iterated for a fixed limit of 50 iterations in the experiments reported here. A tolerance-based stopping rule is also available, terminating when the change in *λ* between successive iterations falls below a user-supplied threshold. On the datasets used here *λ* is stable well before the iteration limit is reached; for the mouse dataset, successive estimates differed by less than 10*^−^*^4^ after approximately 15 iterations.

## Supplementary information

Supplementary Figures,Tables and Notes are provided in a separate Supplementary Information file.

## Data availability

All sequencing datasets and genome assemblies used in this study are publicly available and listed in Supplementary Tables 1 and 3.

## Code availability

kRISP-meR is available at https://github.com/mahmudhera/kRISP-mER.

## References

[1] Doudna, J.A., Charpentier, E.: The new frontier of genome engineering with CRISPR-Cas9. Science 346(6213), 1258096 (2014)

[2] Hsu, P.D., Lander, E.S., Zhang, F.: Development and applications of CRISPR-Cas9 for genome engineering. Cell 157(6), 1262–1278 (2014)

[3] Doudna, J.A.: The promise and challenge of therapeutic genome editing. Nature 578(7794), 229–236 (2020)

[4] Frangoul, H., Altshuler, D., Cappellini, M.D., et al.: CRISPR-Cas9 gene editing for sickle cell disease and *β*-thalassemia. New England Journal of Medicine 384(3), 252–260 (2021)

[5] Gillmore, J.D., Gane, E., Taubel, J., et al.: CRISPR-Cas9 in vivo gene editing for transthyretin amyloidosis. New England Journal of Medicine 385(6), 493–502 (2021)

[6] Kempton, H.R., Qi, L.S.: When genome editing goes off-target. Science 364(6437), 234–236 (2019) 10.1126/science.aax1827

[7] Kleinstiver, B.P., et al.: High-fidelity CRISPR–Cas9 nucleases with no detectable genome-wide off-target effects. Nature 529(7587), 490–495 (2016)

[8] Perez, A.R., Pritykin, Y., Vidigal, J.A., Chhangawala, S., Zamparo, L., Leslie, C.S., Ventura, A.: GuideScan software for improved single and paired CRISPR guide RNA design. Nature Biotechnology 35(4), 347–349 (2017) 10.1038/nbt.3804

[9] Li, T., Wang, S., Luo, F., Wu, F.-X., Wang, J.: MultiGuideScan: a multi--processing tool for designing CRISPR guide RNA libraries. Bioinformatics 36(3), 920–921 (2020) 10.1093/bioinformatics/btz616

[10] Schmidt, H., Zhang, M., Chakarov, D., Bansal, V., Mourelatos, H., Sánchez-Rivera, F.J., Lowe, S.W., Ventura, A., Leslie, C.S., Pritykin, Y.: Genome-wide CRISPR guide RNA design and specificity analysis with GuideScan2. Genome Biology 26(1), 41 (2025) 10.1186/s13059-025-03488-8

[11] Concordet, J.-P., Haeussler, M.: CRISPOR: intuitive guide selection for CRISPR/Cas9 genome editing experiments and screens. Nucleic Acids Research 46(W1), 242–245 (2018) 10.1093/nar/gky354

[12] Doench, J.G., Fusi, N., Sullender, M., Hegde, M., Vaimberg, E.W., Donovan, K.F., Smith, I., Tothova, Z., Wilen, C., Orchard, R., Virgin, H.W., Listgarten, J., Root, D.E.: Optimized sgRNA design to maximize activity and minimize off-target effects of CRISPR-Cas9. Nature Biotechnology 34(2), 184–191 (2016) 10.1038/nbt.3437

[13] Haeussler, M., Schönig, K., Eckert, H., Eschstruth, A., Mianné, J., Renaud, J.-B., Schneider-Maunoury, S., Shkumatava, A., Teboul, L., Kent, J., et al.: Evaluation of off-target and on-target scoring algorithms and integration into the guide rna selection tool crispor. Genome biology 17(1), 148 (2016)

[14] Sun, J., Liu, H., Liu, J., Cheng, S., Peng, Y., Zhang, Q., Yan, J., Liu, H.-J., Chen, L.-L.: CRISPR-Local: a local single-guide RNA (sgRNA) design tool for non-reference plant genomes. Bioinformatics 35(14), 2501–2503 (2019) 10.1093/bioinformatics/bty970

[15] Nurk, S., Koren, S., Rhie, A., et al.: The complete sequence of a human genome. Science 376(6588), 44–53 (2022)

[16] Consortium, H.P.R.: A draft human pangenome reference. Nature 617, 312–324 (2023)

[17] Vollger, M.R., Dishuck, P.C., Sorensen, M., Welch, A.E., Dang, V., Dougherty, M.L., Graves-Lindsay, T.A., Wilson, R.K., Chaisson, M.J.P., Eichler, E.E.: Long-read sequence and assembly of segmental duplications. Nature Methods 16(1), 88–94 (2019) 10.1038/s41592-018-0236-3

[18] Keough, K.C., et al.: AlleleAnalyzer: a tool for personalized and allele-specific sgRNA design. Genome biology 20(1), 1–9 (2019)

[19] Jacquin, A.L.S., Odom, D.T., Lukk, M.: Crisflash: open-source software to generate CRISPR guide RNAs against genomes annotated with individual variation. Bioinformatics 35(16), 3146–3147 (2019) 10.1093/bioinformatics/btz019

[20] Kumbara, A., Tognon, M., Carone, G., Fontanesi, A., Bombieri, N., Giugno, R., Pinello, L.: CRISPR-HAWK: Haplotype- and variant-aware guide design toolkit for CRISPR-Cas. bioRxiv (2025) 10.64898/2025.12.27.696698

[21] Dempster, A.P., Laird, N.M., Rubin, D.B.: Maximum likelihood from incomplete data via the EM algorithm. Journal of the royal statistical society: series B (methodological) 39(1), 1–22 (1977)

[22] Langmead, B., Salzberg, S.L.: Fast gapped-read alignment with Bowtie 2. Nature methods 9(4), 357–359 (2012)

[23] Walker, B.J., Abeel, T., Shea, T., Priest, M., Abouelliel, A., Sakthikumar, S., Cuomo, C.A., Zeng, Q., Wortman, J., Young, S.K., et al.: Pilon: an integrated tool for comprehensive microbial variant detection and genome assembly improvement. PloS one 9(11), 112963 (2014)

[24] Marçais, G., Kingsford, C.: A fast, lock-free approach for efficient parallel counting of occurrences of *k*-mers. Bioinformatics 27(6), 764–770 (2011) 10.1093/bioinformatics/btr011

[25] Engel, S.R., Dietrich, F.S., Fisk, D.G., Binkley, G., Balakrishnan, R., Costanzo, M.C., Dwight, S.S., Hitz, B.C., Karra, K., Nash, R.S., et al.: The reference genome sequence of *Saccharomyces cerevisiae*: then and now. G3: Genes, Genomes, Genetics 4(3), 389–398 (2014) 10.1534/g3.113.008995

[26] Church, D.M., Goodstadt, L., Hillier, L.W., Zody, M.C., Goldstein, S., She, X., Bult, C.J., Agarwala, R., Cherry, J.L., DiCuccio, M., et al.: Lineage-specific biology revealed by a finished genome assembly of the mouse. PLoS Biology 7(5), 1000112 (2009) 10.1371/journal.pbio.1000112

[27] Schneider, V.A., Graves-Lindsay, T., Howe, K., et al.: Evaluation of GRCh38 and de novo haploid genome assemblies demonstrates the enduring quality of the reference assembly. Genome Research 27(5), 849–864 (2017) 10.1101/gr.213611.116

[28] Singh, D., Sternberg, S.H., Fei, J., Doudna, J.A., Ha, T.: Real-time observation of DNA recognition and rejection by the RNA-guided endonuclease Cas9. Nature Communications 7, 12778 (2016) 10.1038/ncomms12778

[29] Singh, D., Wang, Y., Mallon, J., Yang, O., Fei, J., Poddar, A., Ceylan, D., Bailey, S., Ha, T.: Mechanisms of improved specificity of engineered Cas9s revealed by single-molecule FRET analysis. Nature Structural & Molecular Biology 25(4), 347–354 (2018) 10.1038/s41594-018-0051-7

[30] Nurk, S., Koren, S., Rhie, A., Rautiainen, M., Bzikadze, A.V., Mikheenko, A., Vollger, M.R., Altemose, N., Uralsky, L., Gershman, A., et al.: The complete sequence of a human genome. Science 376(6588), 44–53 (2022)

[31] Zhou, Y., Chebotarov, D., Kudrna, D., Llaca, V., Lee, S., Rajasekar, S., Mohammed, N., Al-Bader, N., Sobel-Sorenson, C., Parakkal, P., et al.: A platinum standard pan-genome resource that represents the population structure of asian rice. Scientific data 7(1), 113 (2020)

[32] Shang, L., He, W., Wang, T., Yang, Y., Xu, Q., Zhao, X., Yang, L., Zhang, H., Li, X., Lv, Y., et al.: A complete assembly of the rice nipponbare reference genome. Molecular plant 16(8), 1232–1236 (2023)

[33] Bailey, C.C., Zhong, G., Huang, I.-C., Farzan, M.: Ifitm-family proteins: the cell’s first line of antiviral defense. Annual review of virology 1(1), 261–283 (2014)

[34] Bizard, A.H., Hickson, I.D.: The many lives of type ia topoisomerases. Journal of Biological Chemistry 295(20), 7138–7153 (2020)

[35] Savitzky, A., Golay, M.J.E.: Smoothing and differentiation of data by simplified least squares procedures. Analytical Chemistry 36(8), 1627–1639 (1964) 10.1021/ac60214a047

[36] Rahman, A., Hallgrímsdóttir, I., Eisen, M., Pachter, L.: Association mapping from sequencing reads using k-mers. Elife 7, 32920 (2018)

[37] Vurture, G.W., Sedlazeck, F.J., Nattestad, M., Underwood, C.J., Fang, H., Gurtowski, J., Schatz, M.C.: GenomeScope: fast reference-free genome profiling from short reads. Bioinformatics 33(14), 2202–2204 (2017) 10.1093/bioinformatics/btx153

[38] Ranallo-Benavidez, T.R., Jaron, K.S., Schatz, M.C.: GenomeScope 2.0 and Smudgeplot for reference-free profiling of polyploid genomes. Nature Communications 11(1), 1432 (2020) 10.1038/s41467-020-14998-3

