## Supplementary Information for "kRISP-meR: A Reference-free Guide-RNA Design Tool for CRISPR/Cas9"

### Contents

**1 Supplementary Figures** **2**

**2 Supplementary Tables** **7**

**3 Supplementary Notes** **9**

Supplementary Note 1: Sensitivity to the acceptance threshold . . . . . . . . . . . . . . 9

### 1 Supplementary Figures

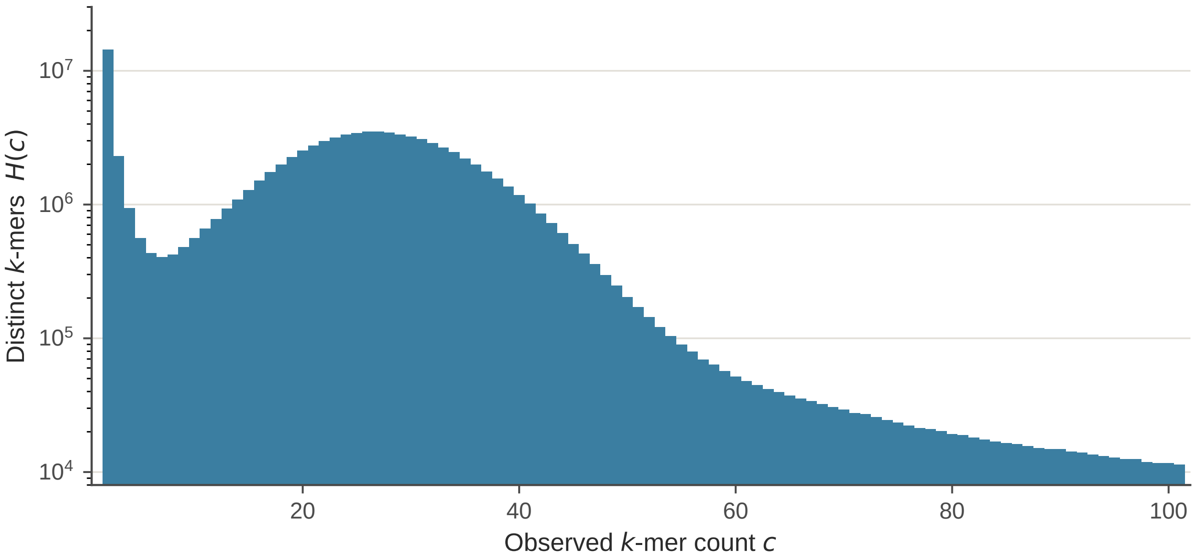


**Supplementary Figure 1:** $k$-mer frequency histogram $H(c)$ for the *H. sapiens* read set, counts $c = 2$ to 101, on a logarithmic vertical scale. The spike at low counts arises from sequencing error. The peak near $c = 26$ corresponds to single-copy genomic $k$-mers and gives the initial estimated $k$-mer coverage $\lambda$. The extended tail reflects sequence present at higher copy number, with components near $2\lambda$, $3\lambda$ and beyond overlapping into it.

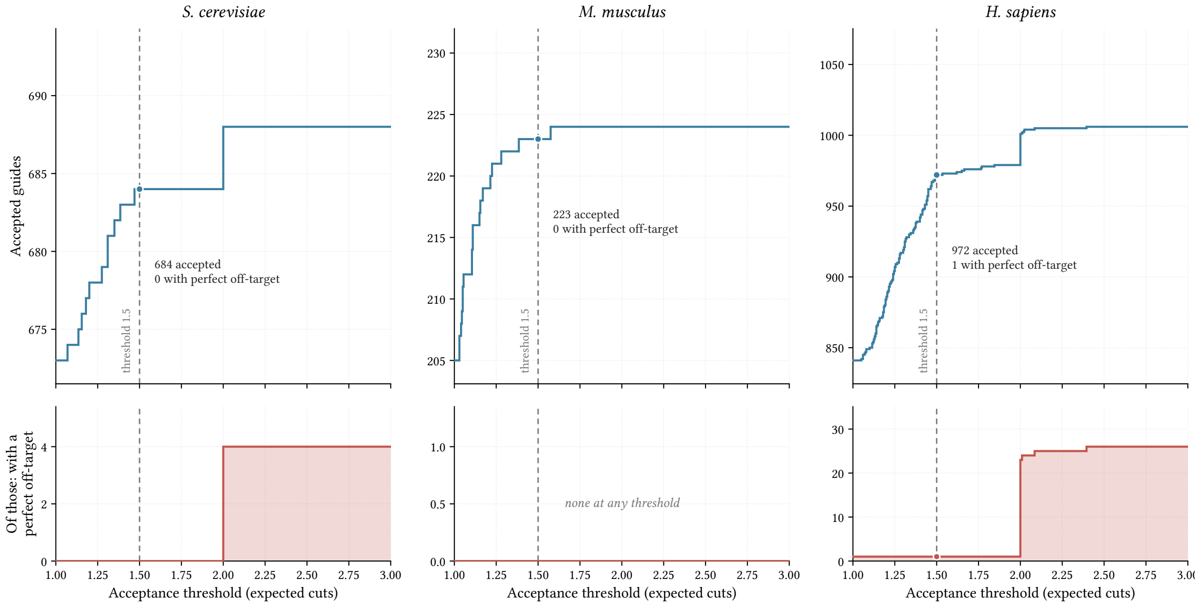


**Supplementary Figure 2:** Effect of the acceptance threshold on kRISP-meR's recommendations. **Top:** number of candidate guides accepted at each threshold on expected cuts. The axis is scaled to the range of the data and does not start at zero. **Bottom:** how many of those accepted guides carry a perfect, zero-mismatch off-target site in the assembly. The dashed line marks the threshold of 1.5 used throughout. See Supplementary Note 1.

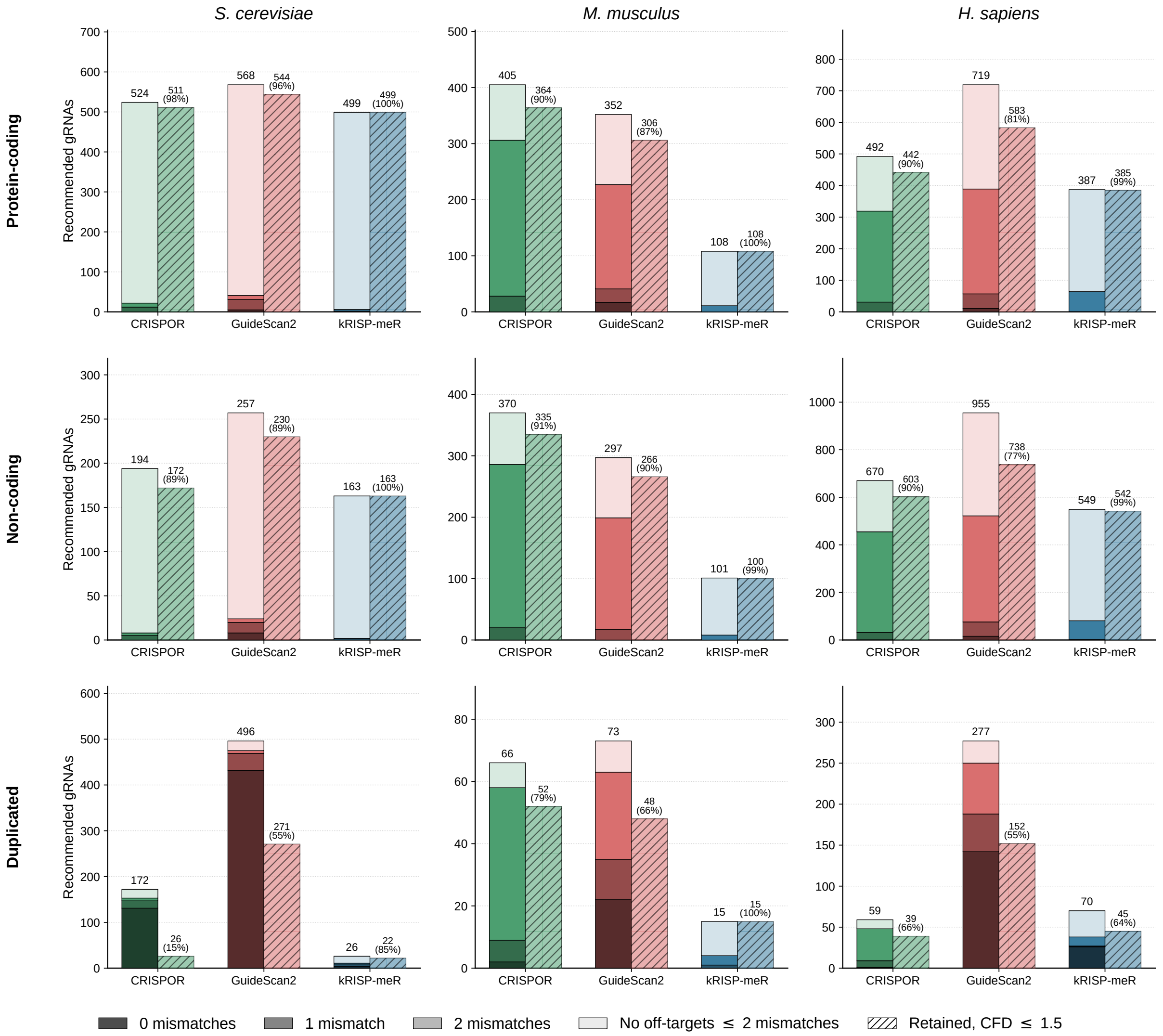


**Supplementary Figure 3:** Off-target profiles of recommended guides by genomic context. Columns are species, in the same order as the main text, and rows are the context from which targets were sampled: protein-coding, non-coding and duplicated regions. In each panel, the solid bar classifies each tool's recommended guides by their most severe off-target site in the reference assembly, with the total above it. The hatched bar gives the number of those guides with assembly-CFD expected number of cuts $\leq 1.5$, with the percentage of the same set. Each guide is counted once, so the three contexts sum to the totals of the off-target breakdown in the main text.

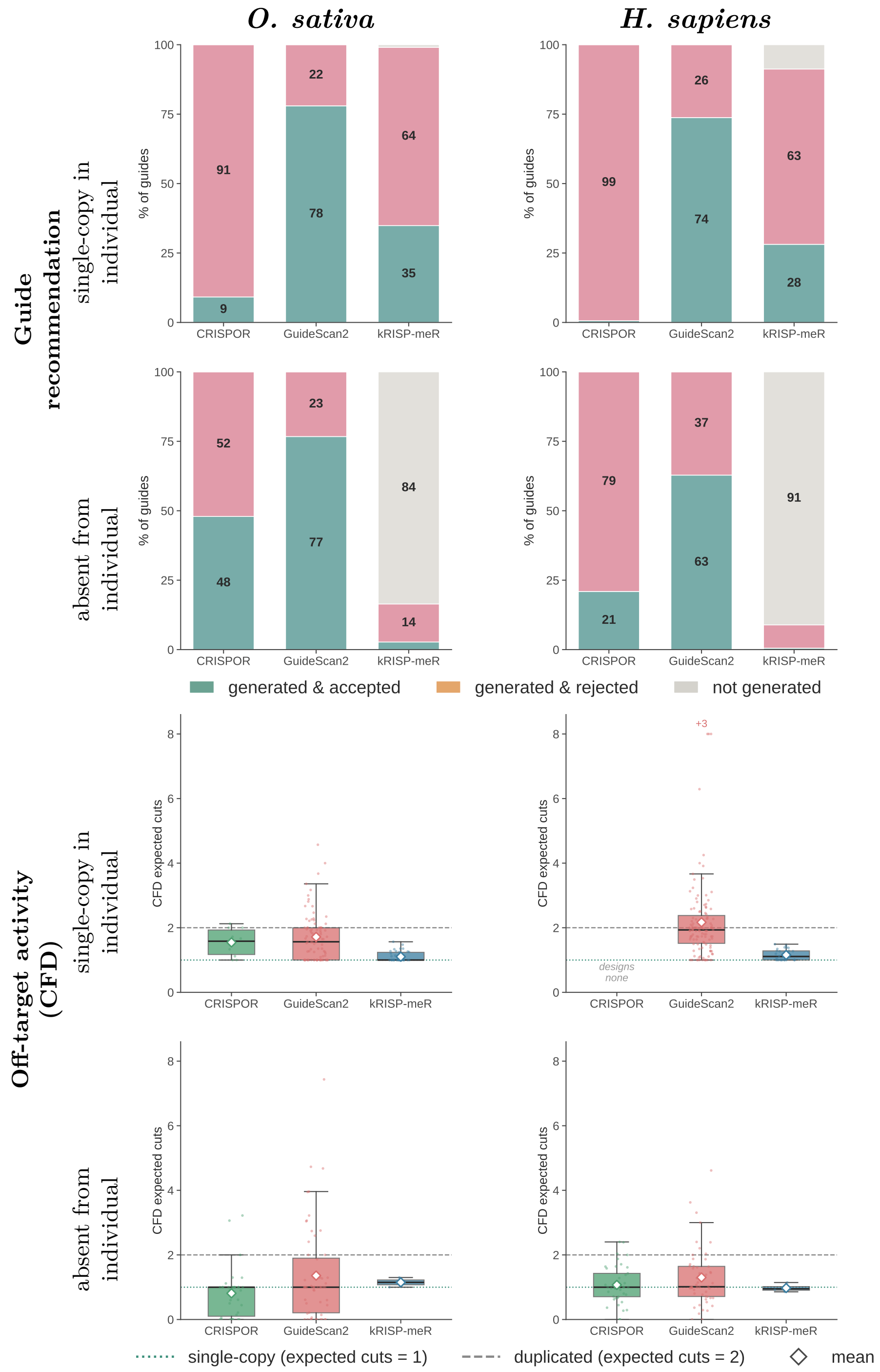


**Supplementary Figure 4:** Guide recommendations and off-target activity for the two discordance categories complementary to those shown in the main text. Rows show loci single-copy in the individual but multiple-copy in the reference, and loci absent from the individual but present in the reference. **Top:** proportion of class-defining guides each tool accepted, rejected, or did not generate. **Bottom:** assembly-CFD expected number of cuts for accepted guides.

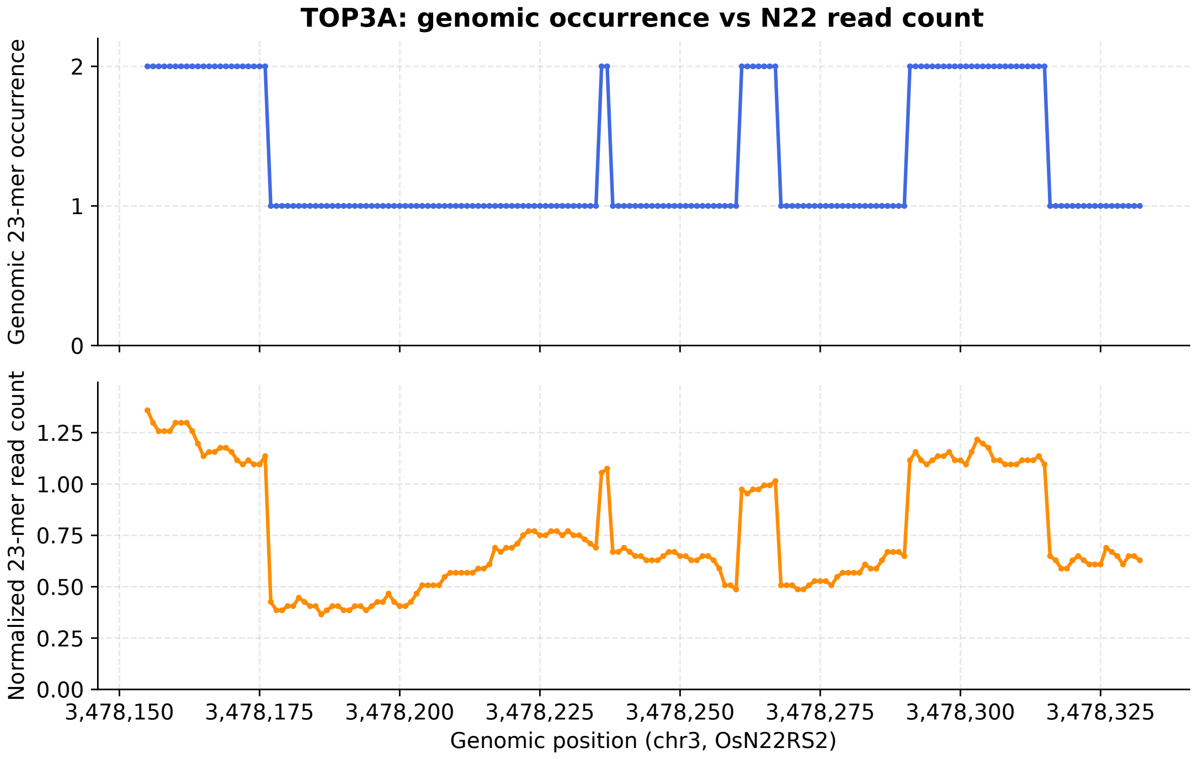


**Supplementary Figure 5:** Occurrence of 23-mers in the assembly and in the reads across the TOP3A target window in *O. sativa* N22 (chr3:3,478,155–3,478,354, OsN22RS2). Each 23-mer is plotted at its lowest coordinate. **Top:** number of occurrences of each 23-mer in the N22 assembly. **Bottom:** count of the same 23-mer in the reads, normalized by the estimated $k$-mer coverage. Read counts step up wherever a 23-mer occurs twice in the assembly.

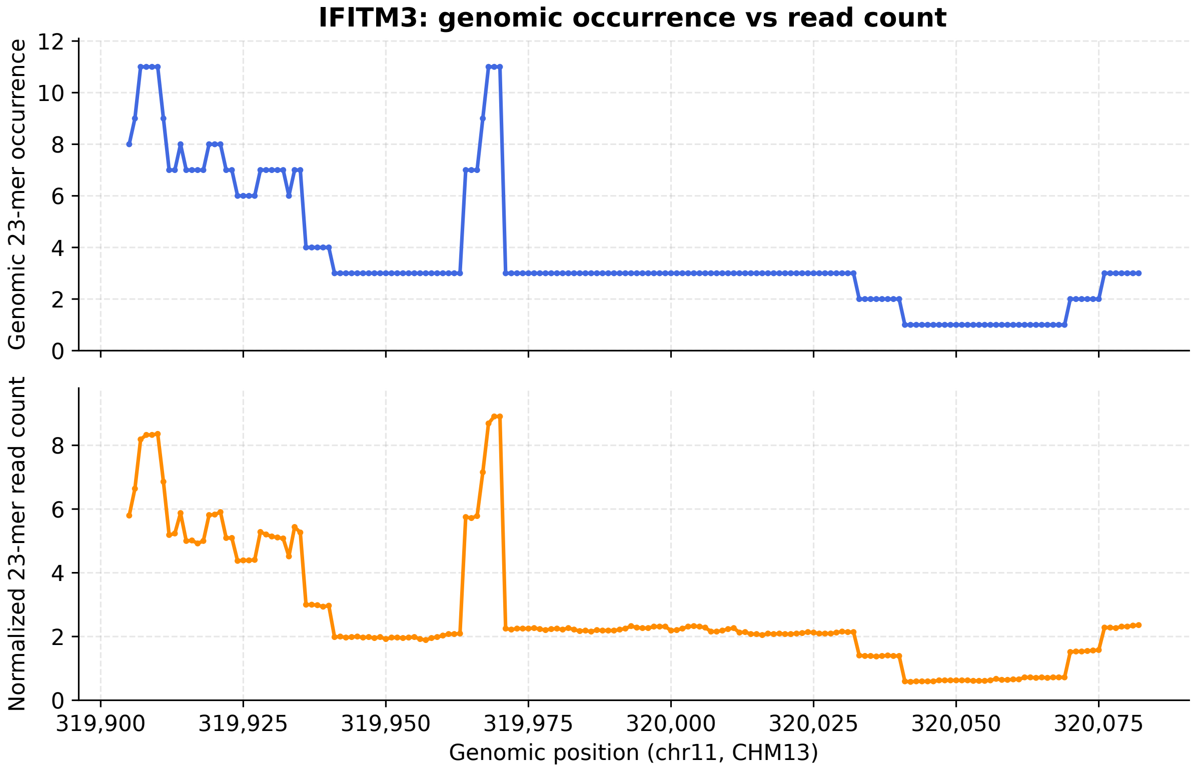


**Supplementary Figure 6:** Occurrence of 23-mers in the assembly and in the reads across the IFITM3 target region in *H. sapiens* (chr11, CHM13v2.0). Panels and plotting position as in Supplementary Figure 5. Read counts follow the same steps, including both peaks, at a lower absolute level. The vertical scales of the two panels differ.

### 2 Supplementary Tables

**Supplementary Table 1:** Sequencing datasets used in the three-species benchmark in the main text. kRISP-meR was run on reads alone; CRISPOR and GuideScan2 were run on the corresponding reference assembly. $\lambda$ is the $k$-mer coverage fitted by the mixture model described in Methods. All datasets are real whole-genome sequencing rather than simulated reads.

| Species | Assembly scored | Reads (SRA) | Read type | $\lambda$ |
|---|---|---|---|---|
| *H. sapiens* | GRCh38.p13 | SRR10035390 (PRJNA559484) | $2 \times 151$ bp | 38.0 |
| *M. musculus* | GRCm39 | SRR3298694 (PRJNA316487) | $2 \times 100$ bp | 18.5 |
| *S. cerevisiae* | S288C (sacCer3) | SRR4244871 | $2 \times 126$ bp | 63.4 |

**Supplementary Table 2:** Counts behind the off-target breakdown of recommended guides in the main text. Recommended guides are classified by their most severe off-target site in the reference assembly. Retained guides are those with assembly-CFD expected number of cuts $\leq 1.5$, computed over the same recommended set.

| | | | Most severe off-target | | | | CFD $\leq 1.5$ | |
|---|---|---|---|---|---|---|---|---|
| Species | Tool | Recommended | 0 mm | 1 mm | $\geq$2 mm | None | $n$ | % |
| *S. cerevisiae* | CRISPOR | 890 | 131 | 33 | 19 | 707 | 709 | 79.7 |
| | GuideScan2 | 1321 | 445 | 75 | 20 | 781 | 1045 | 79.1 |
| | kRISP-meR | 688 | 4 | 9 | 6 | 669 | 684 | 99.4 |
| *M. musculus* | CRISPOR | 841 | 2 | 56 | 592 | 191 | 751 | 89.3 |
| | GuideScan2 | 722 | 39 | 54 | 396 | 233 | 620 | 85.9 |
| | kRISP-meR | 224 | 0 | 1 | 22 | 201 | 223 | 99.6 |
| *H. sapiens* | CRISPOR | 1221 | 2 | 70 | 750 | 399 | 1084 | 88.8 |
| | GuideScan2 | 1951 | 169 | 152 | 840 | 790 | 1473 | 75.5 |
| | kRISP-meR | 1006 | 26 | 4 | 153 | 823 | 972 | 96.6 |

**Supplementary Table 3:** Sequencing datasets and genome assemblies used in the individual–reference genomic-discordance analysis. CHM13 and N22 assemblies were used as operational ground truth to define discordant genomic categories and the candidate-guide pool. kRISP-meR was run only on sequencing reads from the corresponding individual.

| Analysis role | Dataset/assembly |
|---|---|
| *Human* | |
| Reads for kRISP-meR | CHM13, Illumina NovaSeq 6000 whole-genome sequencing, paired-end $2 \times 151$ bp<br>SRR10035390; BioProject PRJNA559484 |
| Individual assembly (operational ground truth) | CHM13v2.0<br>GCA_009914755.4 |
| Reference assembly | GRCh38.p13 primary assembly<br>GCF_000001405.39 |
| *Rice* | |
| Reads for kRISP-meR | *Oryza sativa* cultivar N22<br>SRR4027938, SRR4027939, SRR4027940, SRR4027941 |
| Individual assembly (operational ground truth) | N22, ASM195236v2<br>GCA_001952365.2 |
| Reference assembly | *Oryza sativa* japonica cultivar Nipponbare, ASM3414082v1<br>GCF_034140825.1 |

**Supplementary Table 4:** Guides produced and accepted by each tool on two full-gene targets, and how many accepted guides carry a perfect, zero-mismatch off-target site in the individual's assembly. Guides are counted once by unique sequence. Accepted guides are those with expected number of cuts $\leq 1.5$ (kRISP-meR), specificity score $\geq 60$ (CRISPOR) and specificity $\geq 0.2$ (GuideScan2). Off-target sites were counted from canonical 23-mer counts of the assembly, excluding the on-target site.

| Gene (assembly) | Tool | Produced | Accepted | Accepted with OT | % |
|---|---|---|---|---|---|
| TOP3A (N22) | kRISP-meR | 437 | 145 | 32 | 22.1 |
| | CRISPOR | 818 | 725 | 303 | 41.8 |
| | GuideScan2 | 818 | 770 | 316 | 41.0 |
| IFITM3 (CHM13) | kRISP-meR | 547 | 165 | 0 | 0.0 |
| | CRISPOR | 1174 | 512 | 133 | 26.0 |
| | GuideScan2 | 1168 | 926 | 303 | 32.7 |

### 3 Supplementary Notes

##### Supplementary Note 1: Sensitivity to the acceptance threshold

The acceptance threshold of 1.5 expected cuts governs every comparison in the main text. To examine what this value costs and what it protects against, we counted the guides kRISP-meR accepts at thresholds from 1.0 to 3.0, and how many of those guides carry a perfect, zero-mismatch off-target site in the assembly (Supplementary Figure 2).

Raising the threshold recovers few additional guides. At 1.5, kRISP-meR accepts 684 of 688 candidates in *S. cerevisiae*, 223 of 224 in *M. musculus* and 972 of 1006 in *H. sapiens*. Raising it to 3.0 adds 4, 1 and 34 guides respectively.

Almost every accepted guide with a perfect off-target has an expected-cut value of exactly 2, because a target duplicated in the genome contributes two to the expected number of cuts. At 1.5, no accepted guide carries a perfect off-target in yeast or mouse, and one does in human. At 3.0 the counts rise to 4, 0 and 26. In *H. sapiens*, 22 of the 29 guides gained between 1.5 and 2.0 carry a perfect off-target; in *S. cerevisiae* all 4 of the guides gained do.

A threshold of 1.5 therefore sits below the point at which duplicated targets begin to be accepted, while missing very few usable guides.
